# Hypoxia coordinates the spatial landscape of myeloid cells within glioblastoma to affect outcome

**DOI:** 10.1101/2023.06.30.547190

**Authors:** Michael J. Haley, Leoma Bere, James Minshull, Sokratia Georgaka, Natalia Garcia-Martin, Gareth Howell, David J. Coope, Federico Roncaroli, Andrew King, David Wedge, Stuart Allan, Omar N. Pathmanaban, David Brough, Kevin Couper

## Abstract

Myeloid cells are highly prevalent in glioblastoma (GBM), existing in a spectrum of phenotypic and activation states. We currently have limited knowledge of the tumour microenvironment (TME) determinants that influence the localisation and the functions of the diverse myeloid cell populations in GBM. Here we have utilised orthogonal imaging mass cytometry with single cell and spatial transcriptomics approaches to identify and map the various myeloid populations in the human GBM tumour microenvironment (TME). Our results show that different myeloid populations have distinct and reproducible compartmentalisation patterns in the GBM TME that is driven by tissue hypoxia, regional chemokine signalling, and varied homotypic and heterotypic cellular interactions. We subsequently identified specific tumour sub-regions in GBM, based upon composition of identified myeloid cell populations, that were linked to patient survival. Our results provide new insight into the spatial organisation of myeloid cell sub populations in GBM, and how this is predictive of clinical outcome.

**Teaser:** Multi-modal mapping reveals that the spatial organisation of myeloid cells in glioblastoma impacts disease outcome.

## Introduction

Glioblastoma (GBM), the most common and aggressive type of brain tumour, is almost universally fatal, with a median average survival time of 12-18 months. Despite the tremendous advances in treatment of other cancers, the standard of care treatment for GBM – surgical resection followed by chemoradiotherapy – has remained unchanged for decades *(1)* Indeed, to date, GBM has proven largely refractive to immunotherapies highly effective in other cancers *(2)*. Consequently, there is a critical need to better understand the unique immunobiology of GBMs to develop new effective treatments for this devastating disease.

Myeloid cells (which include monocytes, macrophages and tissue resident microglial cells) are major components of GBM, constituting 30-40% of all cells found within the GBM tumour microenvironment (TME) *(3, 4)*. Myeloid cells are believed to play major roles in promoting GBM progression and treatment resistance, including impairing response to radiotherapy and immunotherapy. As such, they are viewed as attractive targets for novel treatments for GBM, with pre-clinical data supporting the benefit of macrophage modulation *(3)*. However, the myeloid cell compartment is extremely heterogeneous in GBM, being able to adopt a spectrum of pro-inflammatory and suppressive states *(4, 5)* with immunosuppressive phenotypes being associated with worse outcome *(6–11)*.

At present, we have limited knowledge of where the different myeloid cell populations compartmentalise in the complex TME of GBMs, and what tissue factors may shape their positioning and differentiation. This is partly due to the limitations of bulk-based or cell-suspension based methods for studying the GBM TME, which cannot simultaneously capture the spatial and phenotypic heterogeneities seen in myeloid cells in GBM *(4)*. Spatial heterogeneity arises in GBM due to variation in the degree of tumour cell infiltration into healthy tissue, and due to emergence of hallmark features of GBM such as hypoxia-induced necrosis and microvascular proliferation that differentiate it from lower grade tumours *(12)*. Further spatial heterogeneity emerges from the proliferation of specific tumour subclones within different parts of the same tumour *(12, 13)*, and macroscopically in terms of blood flow and tissue perfusion *(14)*. Therefore, there are established spatial features within GBM that may imprint myeloid cell heterogeneity and cell compartmentalisation.

Here, we have combined high-dimensional imaging mass cytometry (IMC) and deconvolution of spatial transcriptomics (ST) datasets to map the myeloid cell compartment in the TME of GBM. Orthogonal validation identified distinct populations of myeloid cells in GBM, with subsets of macrophages and microglia that differed in abundance between areas from the edge and core of the tumour. We found clear compartmentalisation of myeloid populations, with cells of a similar phenotype clustering together (with macrophages showing particularly strong clustering behaviour), and segregating cells of dissimilar phenotypes. The spatial landscape of myeloid cells appeared to be modulated by several putative factors, including cellular interactions (with myeloid, tumour and vascular cells), regional chemokine signalling, and predominantly, tissue hypoxia. Hypoxic niches were preferentially occupied by immunosuppressive macrophages and infiltrating monocytes. Crucially, we identified an association between the transcriptomic signature of specific myeloid cell environments and GBM survival, indicating that the spatial arrangement of specific myeloid cell populations is an important determinant of GBM outcome.

## Results

### Imaging mass cytometry captures the diverse myeloid states present in glioblastoma

Primary IDH-wildtype GBMs were selected from the Department of Cellular Pathology at Salford Royal Hospital (**Table 1**), and edge and core regions of the tumours were annotated by a neuropathologist via histological assessment of H&E stained sections. Tissue micro arrays (TMAs) were subsequently generated (24 × 2.25 mm^2^) and sections were stained with a panel of 37 metal conjugated antibodies, followed by data acquisition on a Hyperion imaging mass cytometer (IMC). Nuclear-based cell segmentation was used to extract the single-cell expression of each marker in the staining panel for downstream analysis (**Fig 1A**).

**Figure 1.**
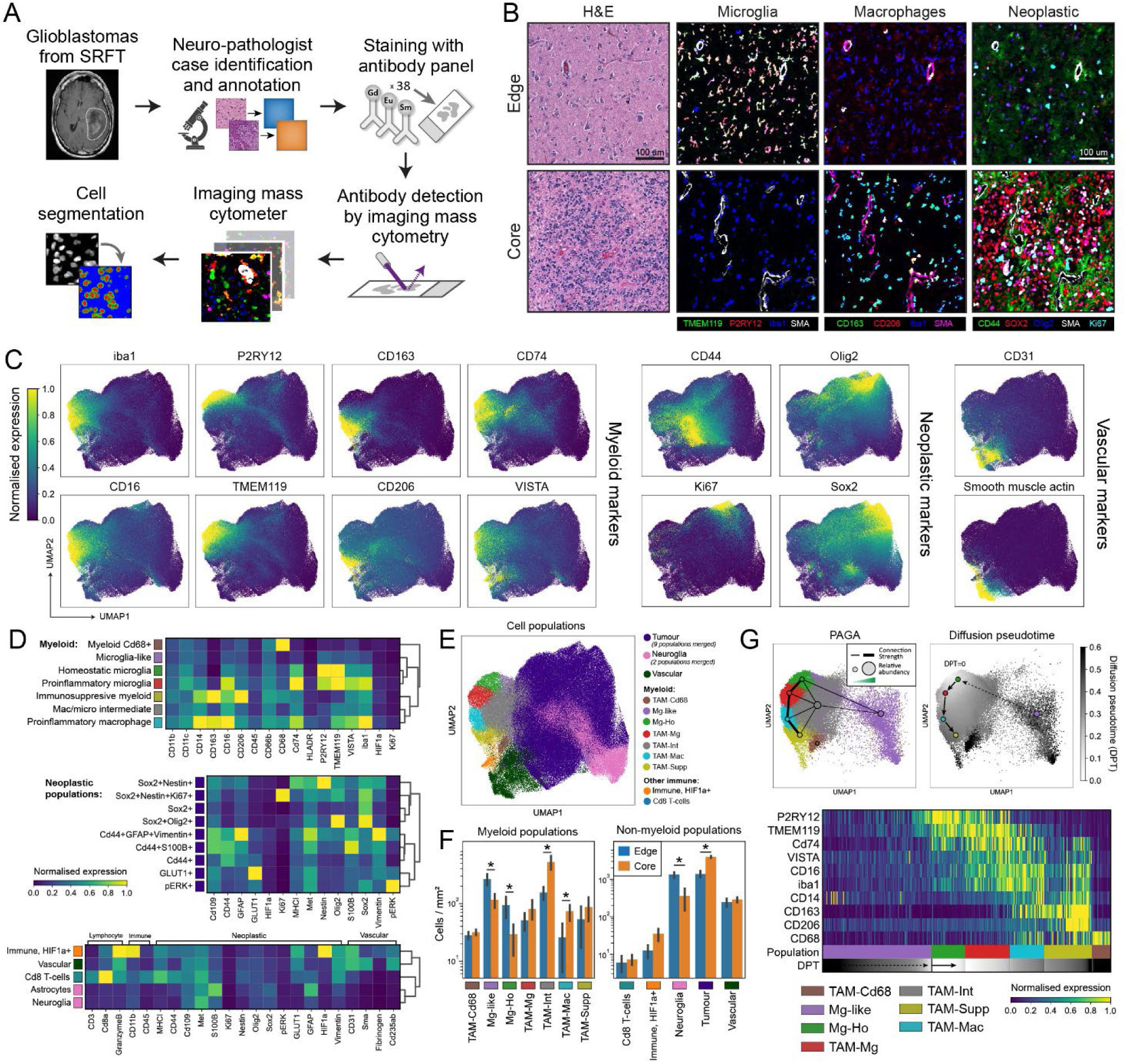
Characterisation of cell populations present in the GBM tumour microenvironment using imaging mass cytometry. **A**. Overview of the glioblastoma patient samples and imaging mass cytometry workflow. **B**. Representative IMC and H&E images from GBM cases taken from either the edge or core of the tumours, visualising key microglial, macrophage and neoplastic markers. **C**. UMAPs visualising the single-cell data acquired from the IMC workflow, demonstrating the distribution of the myeloid, neoplastic, and vascular markers over all cells. Each marker is normalised to the 99^th^ percentile of its expression. **D**. Heatmaps showing the mean marker expression for the populations identified in the single-cell IMC data using Leiden clustering. **E**. UMAP showing the labelled cell populations resulting from Leiden clustering. **F**. Comparison of the abundances of the different myeloid and non-myeloid populations between the edge and core regions of the tumours. *p<0.05, groups compared by multiple linear regression. **G**. Diffusion pseudo time and PAGA analysis demonstrating a pathway of myeloid differentiation starting at homeostatic microglia (Mg-Ho), through to proinflammatory activation microglia (TAM-Mg), proinflammatory macrophages (TAM-Mac), and ultimately to immunosuppressive myeloid cells (TAM-Supp).

**Table 1.** Clinical characteristics of samples for imaging mass cytometry.

| Patient | Site | Age at surgery | Sex | IDH1 | ATRX | MGMT |
| --- | --- | --- | --- | --- | --- | --- |
| 1 | Left temporal | 63 | F | WT | WT | Not performed |
| 2 | Left temporal | 73 | F | WT | WT | Hypermethylated |
| 3 | Right parietal | 63 | F | WT | WT | Unmethylated |
| 4 | Left temporal | 53 | M | WT | WT | Unmethylated |
| 5 | Left parietal | 54 | F | WT | WT | Hypermethylated |
| 6 | Left frontal | 62 | F | WT | WT | Hypermethylated |
| 7 | Left temporal | 73 | M | WT | WT | Unmethylated |
| 8 | Left cingulate | 62 | M | WT | WT | Not performed |

IMC allowed us to simultaneously identify and contrast the components and spatial organisation of the tumour microenvironment within the edge and core regions *in situ*. Given the role of myeloid cells in GBM *(5)*, we designed our antibody panel and subsequent analyses to capture the diversity of myeloid cell populations found within GBM. IMC allowed us to putatively identify microglia (iba1+, TMEM119+, P2RY12+) and tumour associated macrophages (iba1+, CD163+) based upon their morphology and marker expression, as well as cells positive for markers of CNS progenitor cells and cellular proliferation often associated with neoplastic cells (CD44, SOX2, Olig2, Ki67) (**Fig 1B**). Representative images suggested the abundance of microglia and TAMs varied between the edge and core of the tumour, as previously reported *(9, 15–17)*.

To understand the expression patterns of the analysed molecules within the single cells segmented within our dataset, we first visualised their distribution using UMAP. This showed clear segregation between cells expressing myeloid, neoplastic, and vascular makers (**Fig 1C, Supplemental 1A**). Within the region corresponding to myeloid cells, there was separation between cells with highest expression for markers of microglia (TMEM119+, P2RY12+) and tumour associated macrophages (iba1+, CD163+). However, the lack of a discrete boundary between microglia and TAMs cell clusters, and the distribution pattern of other myeloid markers (CD74, CD16, VISTA, CD206), suggested the presence of multiple closely and distantly related myeloid subpopulations in the GBM tumour. Similarly, neoplastic markers were heterogeneously expressed throughout the neoplastic compartment of the UMAP, likely corresponding to distinct neoplastic populations.

To identify the cell populations present in our IMC data, Leiden clustering was performed *(18)*, resulting in profiling of 21 distinct myeloid and non-myeloid populations (**Fig 1D, E**). Within these, seven were identified as myeloid populations, nine populations were identified as tumour cells, two corresponded to neuroglia (astrocytes and other CNS cells), three to non-myeloid immune populations (lymphocytes and a HIF1a+ immune population), and one to vascular cells (**Fig 1E).** Two populations were identifiable as microglia. The first showed higher expression of the homeostatic microglial marker P2RY12 (Mg-Ho; homeostatic microglia), and the second being pro-inflammatory (TAM-Mg; proinflammatory microglia), with lower P2RY12 and increased expression of markers of proinflammatory activation (iba1^high^, VISTA+, CD16+, CD74^high^). Two populations were identified as tumour associated macrophages (CD163+), with one having a proinflammatory phenotype (TAM-Mac; tumour associated macrophages), and moderate expression of TMEM119. The other TAM population had comparatively lower expression of CD14 and CD16, but high expression of CD206 (TAM-Supp; immunosuppressive myeloid), a phenotype which has suggested to reflect immunosuppressive myeloid cells in GBM *(11, 17, 19, 20)*. Besides populations with defined phenotypes, we also found a sizeable population of intermediate TAM/microglial population without a canonical microglia or macrophage signature, as has previously been reported *(21, 22)*. We also found a population of myeloid cells with low expression of myeloid markers relative to other myeloid cells which may represent cells with low activation (Mg-like; microglia-like), and a population which were only positive for CD68 (TAM-Cd68). Although clustering did identify putatively different tumour cell populations, our panel did not robustly distinguish the neoplastic subtypes known to be present in GBM *(23)*. As such, to facilitate downstream analyses, we merged tumour populations into a single population. Cells classified as neuroglia did not express markers that would otherwise identify them as myeloid, tumour or vascular cells, and thus likely represent a mixed population of glia and other CNS-resident cells.

### The abundance of myeloid populations varies in the different GBM regions

Previous studies have reported that microglia and macrophages are preferentially found at the periphery of the tumour, or the core of the tumour, respectively *(9, 15–17)*. We therefore calculated the abundance of the 7 myeloid subpopulations, and non-myeloid cells, within the histologically validated edge and core regions of the tumour (**Fig 1F**). We found a significant increase in the numbers of homeostatic microglia and microglial-like cells in the edge, and an increase in intermediate and proinflammatory populations (TAM-Int, TAM-Mac) in the core. Hierarchical clustering of regions, based upon relative myeloid abundances, further indicated that edge and core regions contained distinct myeloid cell compartments (**Supplemental 1B**). We also saw an increase in neuroglial cells in the tumour edge, and, as expected, an increase in tumour cells in the core. Correlating the abundances of the various populations within each region of interest showed distinct clusters of positively and negatively correlated populations (**Supplemental 1C**). For example, microglia (Mg-Ho and Mg-like) and neuroglial cells abundances correlated positively with each other, but negatively correlated with TAM populations (TAM-Mac, Tam-Supp) and tumour cell numbers. Notably, although there were differences in abundances of myeloid subpopulations between core and edge regions, most populations were found in all regions analysed (**Fig 1F**, **Supplemental 1D)**. Together, these analyses suggest that although there may be broader features of the TME that promote the accumulation of microglia versus tumour associated macrophages, there may be more specific local features present in regions which drive the accumulation of specific myeloid subpopulations.

### Myeloid populations exist on a spectrum of differentiation and activation states in GBM

Our IMC data, along with published single-cell sequencing studies, demonstrate that myeloid cells exist in a spectrum of activation states in GBM *(7, 17)* with the polarisation of myeloid cells towards either microglia or TAMs being the most prominent *(19, 24, 25)*. However, the exact ontogeny of the identified myeloid populations in GBM remains unclear *(4)*. We therefore used PAGA analysis to understand the relationships between the 7 myeloid populations identified by IMC (**Fig 1G**). This showed strong connectivity, and therefore a pathway of differentiation, between 4 myeloid populations: homeostatic microglia (Mg-Ho), proinflammatory microglia (TAM-Mg), proinflammatory macrophages (TAM-Mac), and immunosuppressive myeloid cells (TAM-Supp). As microglia (Mg-Ho) are the brain resident immune cells, we then assessed the differentiation from this native state using diffusion pseudotime analysis. Plotting pseudotime against the route of differentiation defined by PAGA demonstrated that microglial markers are lost in favour of activation markers (VISTA, CD74, CD16, iba1 upregulation), and eventually transitioning into an immunosuppressive phenotype characterised by upregulation of CD163 and CD206 *(17, 20)*. Together, these analyses suggest endogenous microglia may differentiate into tumour associated macrophages, and that the primary pathway through which this occurs is via a state of proinflammatory activation. An alternate pathway by which cells first transition through intermediary state (TAM-Int) before differentiating into immunosuppressive TAMs is less well supported by the PAGA and pseudotime analyses (**Supplemental 1E**).

### Microglia and macrophages show conserved patterns of compartmentalisation in GBM

Although we observed different abundances of myeloid populations between core and edge regions, our data clearly identified that multiple different myeloid cell populations were present within the same areas of the tumour. This raised the question whether myeloid cell compartmentalisation in GBM is stochastic, or whether there are deterministic cellular and environmental drivers of myeloid positioning and function. To investigate this, the previously identified myeloid populations (**Fig 1D**) were mapped back to their locations in the tumour (**Fig 2A**). This revealed that different myeloid populations had different distributions in the TME, with some populations appearing to cluster in specific areas (e.g. TAM-Supp), others aggregated more loosely (e.g. TAM-Int) whereas others appeared more evenly distributed in the TME (e.g. Mg-Ho). Qualitatively, these patterns seemed conserved between the core and edge of the tumour. Notably, myeloid cells appeared to spatially associate with other myeloid cells in a similar position on the microglial-TAM differentiation axis defined by diffusion pseudotime (**Fig 2B**, pseudotime defined in **Fig 1G**).

**Figure 2.**
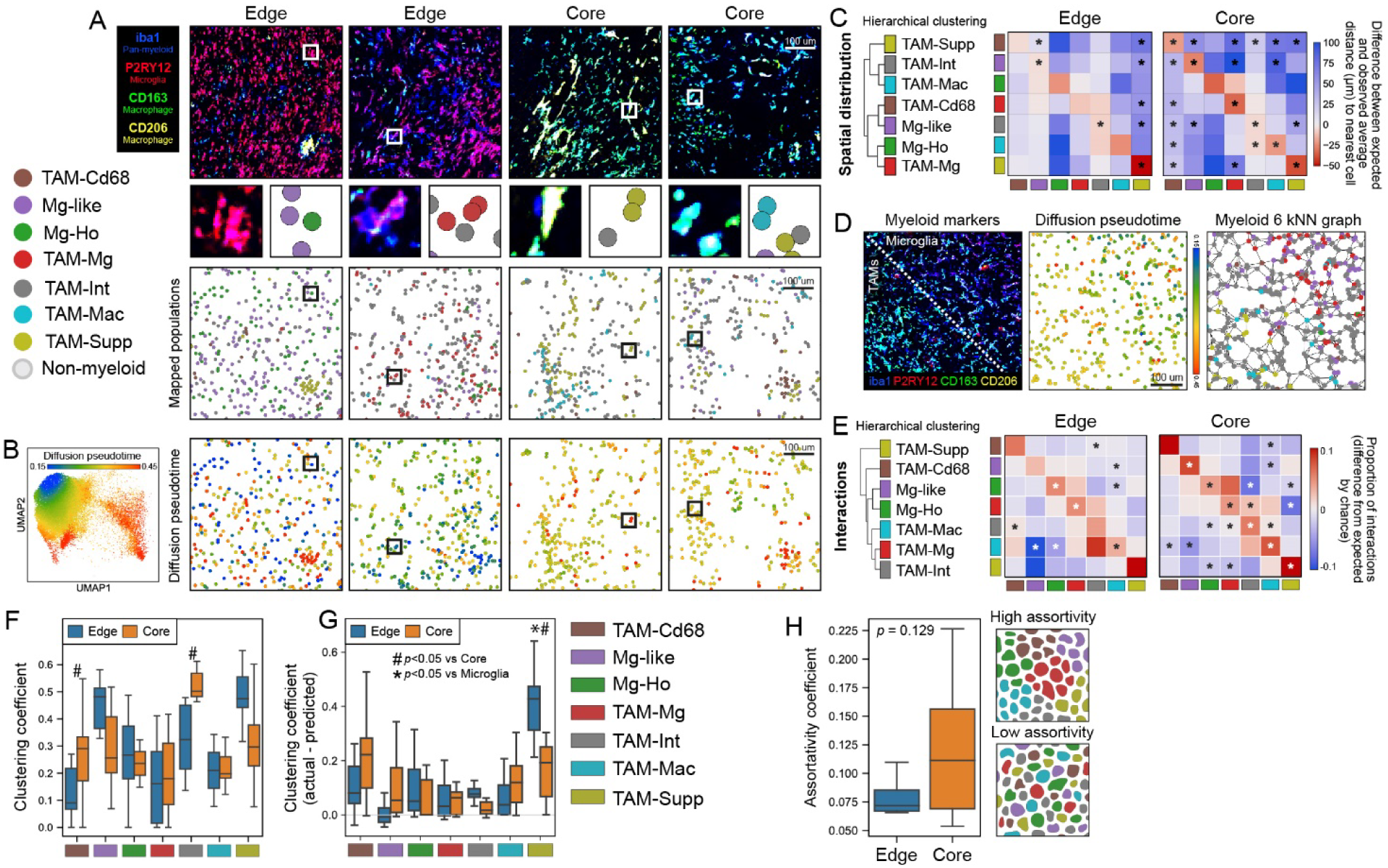
Myeloid cells exhibit high homotypic and low heterotypic clustering behaviour in GBM. **A**. Mapping of the myeloid populations identified from single-cell analyses to their locations in the TME in edge and core regions. **B**. Diffusion pseudotime showing how differentiation away from a homeostatic microglial phenotype relates to myeloid cell positioning. **C**. Spatial distribution analysis which shows whether populations are closer or further away from each other in the TME than would be expect by chance. **D**. IMC image showing separation of microglia (P2RY12+) and macrophages (CD163+, CD206+), alongside the position of the cells on the microglia-macrophage diffusion pseudotime axis, and how the cells are connected in the 6 nearest neighbours analyses. **E**. Proportion of cellular interactions made by myeloid populations when each myeloid cell is connected to its 6 nearest neighbours. Hierarchical clustering of the interactions shows that cells with a similar phenotype also have similar proportions of interactions with other populations. **F**. Clustering coefficient of the different populations in the edge and core of the tumour. **G**. Correction of clustering coefficients for differences in cell abundance, in which a positive value suggests cells are clustering at a greater rate than expected by chance. **H**. Comparison of assortativity of myeloid populations in edge and core regions. Comparison made by Wilcoxon test with Benjamini-Hochberg correction (**C, E**), multiple linear regressions, corrected for multiple comparisons by Holm-Šídák (**F, G**), or Mann-Whitney U test (**H**)

To prove whether there was a preference for myeloid populations to localise close to or avoid other populations in the tumour microenvironment, we quantified whether the observed distance between myeloid populations was significantly different to the distance predicted by a random distribution of cells (**Fig 2C)**. Hierarchical clustering of the resulting data demonstrated that microglial (Mg-like, Mg-Ho, TAM-Mg) and macrophage subpopulations (TAM-Mac, TAM-Supp) have distinct patterns of spatial distribution. Specifically, microglial and macrophage populations appeared to show a preference for spatial segregation, with macrophage populations associating with one-another and localising away from microglia, and microglia enriching with other microglia and localising away from macrophages. All populations had the greatest spatial enrichment with themselves. In some populations (TAM-Cd68, TAM-Int, Mg-like), the difference between the expected and observed distances to other populations was closer to zero, though often still significant. This suggests that these populations are closer to a stochastic distribution, being more randomly and evenly distributed throughout the TME. As the overall pattern of interactions was broadly similar between edge and core regions, it suggests a conservation of the propensity of myeloid cells with a similar phenotype to spatially associate together even in the context of different cell abundances and environments throughout the tumour. To assess whether the spatially associated myeloid populations were interacting in the GBM TME, myeloid cells in each region were subsequently expressed as networks, where each cell was connected to its 6 nearest neighbours (**Fig 2D**, **Supplemental 2A**). This analysis supported our previous observations, showing that myeloid populations most significantly interact with themselves, interact to some degree with phenotypically similar populations, and tend not to interact with dissimilar populations (**Fig 2D&E)**. As with previous analyses, this pattern of behaviour was similar between edge and core regions, suggesting this behaviour is conserved throughout the tumour.

The tendency of myeloid cells to spatially associate with cells of the same type was investigated by measuring each populations’ clustering coefficient (**Fig 2F**). Clustering coefficient is a descriptive statistic of network properties in which a high value designates that a population forms densely interconnected clusters *(26)*, and a low value suggests cells of that population are weakly connected and more loosely clustered in the TME). In this initial analysis, all myeloid populations showed a similar propensity for clustering, with significantly higher clustering in TAM-Cd68 and TAM-Int populations in the core. These clustering values suggest that most myeloid populations form small, weakly connected clusters in the TME. As more abundant populations could be clustering purely by chance, we then corrected clustering coefficients for differences in population abundance between regions. This showed that almost all populations showed more clustering than would be expected by chance (**Fig 2G**). Notably, TAM-Supp cells showed significantly more dense clustering in the edge. The pattern of clustering was otherwise conserved between edge and core regions. Measuring the assortativity (a descriptive statistic of the tendency of populations in a network to connect to populations of the same type over different populations *(27)*) similarly suggested that cells showed a weak but positive preference to connect to cells of the same population, and that this was similar between edge and core regions (**Fig 2H**). Overall, these data demonstrate that different myeloid populations segregate and form loose homotypic clusters in the TME, and that the biological drivers of this segregation are mostly independent of broader location in the edge or core of the tumour.

### The positioning of myeloid cell populations is impacted by tumour, neuroglial and vascular interactions in the GBM TME

Our data suggest myeloid cells are not randomly distributed in the GBM TME. We hypothesised that this myeloid compartmentalisation could be driven by myeloid cells interacting with other highly abundant nonmyeloid cells in the TME, such as tumour (SOX2), vascular (CD31, SMA), and neuroglial cells (defined in **Fig 1)**. Indeed, when we mapped the myeloid populations alongside these other cell types in the edge and core, low activation microglial populations (Mg-like, Mg-Ho) were qualitatively associated with neuroglial cells, TAM-Supp cells were often associated with vascular cells, and proinflammatory populations (TAM-Mg, TAM-Mac) were associated with tumour cells (**Fig 3A**).

**Figure 3.**
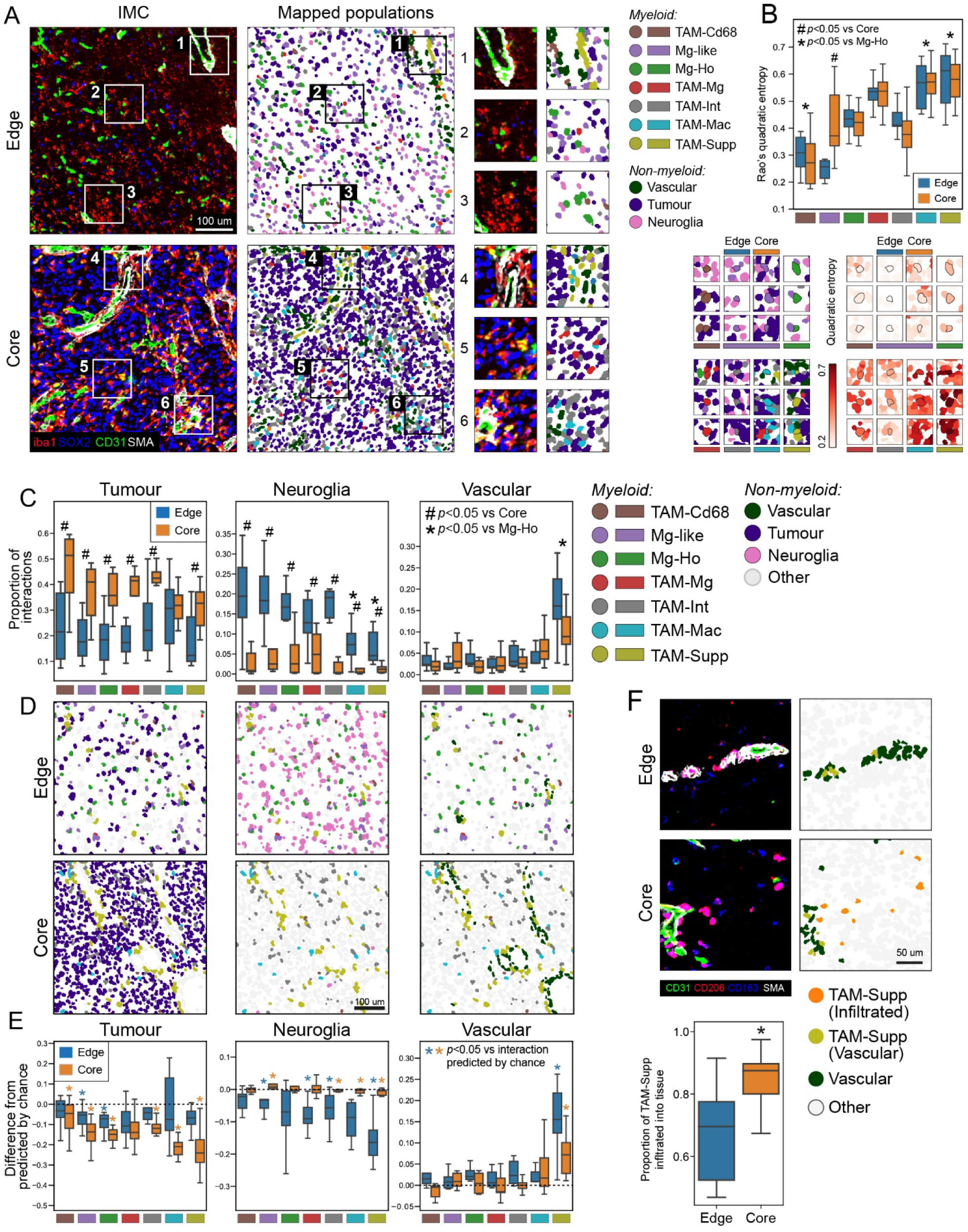
Myeloid cell interactions with tumour, neuroglial and vascular cells in GBM. **A**. Myeloid cells mapped alongside tumour, neuroglial and vascular cells in representative edge and core regions. **B**. Quadratic entropy for the myeloid populations in the edge and core of the tumour. This quantifies how heterogeneous a cell is with respect to its interacting cell, with high values indicating a cell interacts with several cells with different phenotypes. **C**. Proportion of direct cell-to-cell interactions made by myeloid populations with non-myeloid cells, compared between edge and core regions. **D**. Representative examples of myeloid populations mapped alongside non-myeloid populations. **E**. Difference between the observed rate of cell-to-cell interaction data (see **C**), and the rate of interactions predicted by chance. This analysis corrects for the established differences in abundances of the myeloid and non-myeloid cells between different regions. **F**. Comparison between edge and core regions in the rate of infiltration of TAM-Supp cells into the parenchyma, defined here as being 10 µm away from nearest vascular cell. Comparisons made by linear mixed models (**B**, **C**) or Wilcoxon tests (**E**) with Holm-Šídák corrections, or Mann-Whitney U test (**F**).

To quantify the different environments in which the different myeloid cell populations were found we used a local measure of Rao’s quadratic entropy (**Fig 3B**). This measures the phenotypic diversity between each myeloid cell and the cells (non-myeloid and myeloid) with which it interacts, with highly scoring cells interacting with several cells with heterogeneous phenotypes. This demonstrated that macrophage populations (TAM-Mac, TAM-Supp) had significantly greater quadratic entropy than microglia. Whilst we have previously established that myeloid populations cluster in the tissue (**Fig 2**), these analyses further suggest that macrophages form clusters in more cellularly heterogenous areas of the tumour, interacting with multiple non-myeloid cells with diverse phenotypes. In comparison, microglia have less diversity in their interactions with other non-myeloid cells. The only difference between edge and core was in Mg-like cells, where cells in the core had significantly greater quadratic entropy. This suggests that the local cellular organisation of non-myeloid cells influences myeloid cell localisation in a way that is conserved throughout the tumour, so that macrophages preferentially cluster in cellularly dense and phenotypically diverse areas of the tumour.

Quantification interactions between myeloid cells and tumour, neuroglial and vascular cells demonstrated that almost all myeloid populations had significantly greater proportion of their cellular interactions with tumour cells in the core, and significantly greater interactions with neuroglial cells in the edge. TAM-Mac and TAM-Supp populations also had significantly less interactions with neuroglial cells compared to microglia (**Fig 3C**). Notably, the greatest proportion of myeloid cell interactions were with other myeloid cells in both the core and edge, supporting our earlier observations of myeloid cell clustering and segregation (**Supplemental 2B**). The only myeloid population to show significant interaction with vascular cells was the TAM-Supp population, which was enriched in both edge and core. Whilst this suggested there were distinct neoplastic and neuroglial influence on myeloid cell positioning in edge and core, these effects could be impacted by established differences in myeloid abundance in different regions and cases (**Fig 1F**, **Fig 3D**). For example, highly abundant populations may be co-localising, and so appear to be interacting, purely by chance. Once we corrected rates of cellular interactions for differences in cellular abundances in the edge and core regions, myeloid populations were found to avoid interacting with tumour cells equally in both core and edge regions, with most populations showing significantly less interactions than would be expected by chance (**Fig 3E**).

TAM-Mac and TAM-Supp cells showed the greatest avoidance of tumour cells, particularly in the core. Furthermore, abundance-corrected interactions suggested myeloid populations also avoided interacting with neuroglial cells, particularly in the edge, though also to a lesser extent in the core. The greatest avoidance of neuroglial cells was seen in TAM-Supp cells in the edge. In corrected values, TAM-Supp cells continued to be the only population with a significant interaction with vascular cells in both edge and core cases, but differences in the strength of association was identified in the two different tumour regions (**Fig 3E**). As we also previously observed differences in the clustering behaviour of TAM-Supp cells between edge and core regions (**Fig 2F**), we subsequently calculated the proportion of TAM-Supp cells that had infiltrated into the tissue in the edge and core regions of the tumour. This showed that significantly more TAM-Supp cells had infiltrated into the tissue in the core, suggesting the vasculature may be a point of entry for TAM-Supp cells, which remain primarily associated with the vasculature in edge regions, but infiltrate into the tumour in the core (**Fig 3E**)

Overall, myeloid cells showed a propensity to avoid interacting with tumour and neuroglial cells that was broadly similar between different myeloid populations. Observations were also similar between edge and core, other than neuroglial cells having reduced impact on myeloid positioning in core regions. This suggests that differences in localisation of specific myeloid populations is not strongly driven by the avoidance of (or preference for) direct heterotypic cell-to-cell interactions with neuroglial or tumour cells. The observed co-localisation of macrophages with tumour cells may therefore be driven by common factors or tissue signals that do not rely on cell-cell interactions. By comparison, there was a clear population-specific association of TAM-Supp cells with the vasculature.

### Macrophages preferentially localise to hypoxic areas of the TME in GBM

Previous analyses suggested cell-intrinsic behaviours associated with specific myeloid cell populations, or common to all GBM-associated myeloid populations, are at least partly responsible for myeloid positioning in the TME. However, environmental factors and biological niches likely also affect myeloid cell activities and positioning. Tissue hypoxia, often leading to necrosis, is a defining feature of glioblastoma. We therefore hypothesised that compartmentalisation of different myeloid cell populations may be driven by the degree of hypoxia in the surrounding tissue, assessed here by an upregulation of GLUT1 and pERK, markers which have been previously shown to increase in response to tissue hypoxia (Muz 2015). Within the regions analysed, markers of hypoxia (GLUT1, pERK) were not uniformly distributed, suggestive of discrete hypoxic niches within the tumour (**Fig 4A**). Specifically, GLUT1 was constrained to the vasculature in edge cases, but was found more diffusely in the parenchyma in the core in a pattern which has previously been observed in GBM and is thought to represent metabolic adaptation to tissue hypoxia *(28)*. We therefore measured the GLUT1 and pERK staining in the local environment surrounding each myeloid cell (**Fig 4B, C**). This demonstrated that macrophage populations (TAM-Mac, TAM-Supp) were in environments with significantly higher expression of GLUT1 and pERK than microglia, with expression being highest in the TAM-Supp population. This suggests that macrophages preferentially localise to areas of hypoxia, whilst microglia are typically found in comparatively normoxic areas. However, vascular-associated GLUT1 may also be contributing to the association of TAM-Supp cells with GLUT1, as we have previously established they cluster in vascular locations (**Fig 3**).

**Figure 4.**
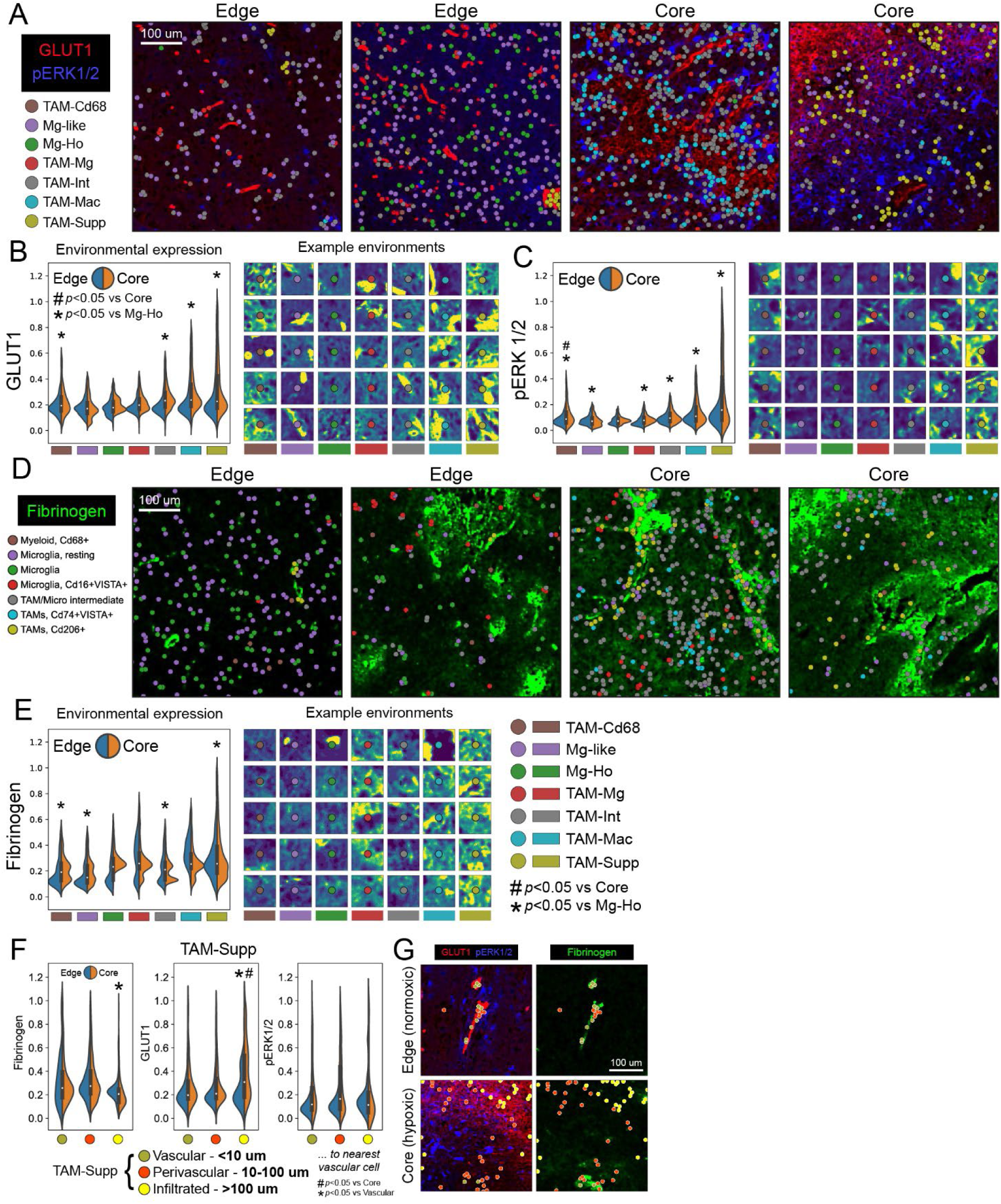
Association of hypoxia and fibrinogen with myeloid cell positioning in GBM. Myeloid populations mapped alongside markers of (**A**) environmental hypoxia in GLUT1 and pERK1/2 staining, or (**D**) fibrinogen. Quantification of environmental GLUT1 (**B**), pERK1/2 (**C**), and fibrinogen (**E**) staining around each myeloid population in edge and core regions. A cell’s environment was defined as a 40 µm square centred on the cell. **F**. Comparison of environmental stains in TAM-Supp cells that were either vascular associated, perivascular, or fully infiltrated into the tumour, in both edge and core regions. **G**. Representative examples of TAM-Supp cells with different vascular associations in a normoxic edge region, and a hypoxic core region. Comparisons made by linear mixed models with Holm-Šídák corrections (**B**, **C**, **E, F**). A further environmental factor and feature of GBM pathology is break down of the blood-brain barrier (BBB), which is associated with inflammation and aberrant angiogenesis *(14)*. We therefore identified areas of BBB damage in both the edge and core of the tumour by measuring the presence of fibrinogen in the tumour, which was restricted to the lumen in vessels with an intact BBB, but which leaked into the brain when the BBB was compromised (**Fig 4D**). When we quantified the environmental localisation of fibrinogen around each cell, we found that TAM-Supp cells localised to areas with significantly higher fibrinogen than microglia (**Fig 4E**). This suggests that TAM-Supp cells localise to areas of BBB breakdown. However, as we previously established that TAM-Supp cells are vascular-associated, fibrinogen trapped in intact vessels could contribute to this environmental enrichment. We therefore repeated previous analyses but separated TAM-Supp cells into vascular, perivascular, or fully infiltrated. This demonstrated that TAM-Supp cells in vascular and perivascular locations had similar environmental localisation of markers, but that cells that had fully infiltrated had significantly lower fibrinogen but higher GLUT1 association (**Fig 4F&G**). Although most TAM-Supp cells were vascular associated (**Fig 3**), these results suggest that infiltrative TAM-Supp cells localise in areas of hypoxia, and thus that hypoxia may be a signal that draws them away from their usual vascular and perivascular locations.

### Myeloid populations identified by IMC in GBM align with those defined through single-cell RNA sequencing (scRNAseq)

We next took an orthogonal approach to validate the environmental drivers of myeloid cell positioning. We characterised the myeloid cell populations present in GBM tumours by re-analysing a published scRNAseq dataset *(29)*. This identified six populations, including two populations of microglia (Mg-Ho; homeostatic microglia, and TAM-Mg; proinflammatory microglia), two of macrophage-derived TAMs (TAM-Mac and TAM-Mac-Supp), a population of TAM-microglial intermediate cells (possessing features of both microglia and macrophages), and monocytes (**Fig 5A&B, Supp 3A**). Clear differences were found in inflammatory, metabolic, and proliferative signalling between these populations (**Supp 3B & C**). Importantly, these populations aligned with IMC populations, showing a similar distribution and polarisation of markers between modalities (**Supp 3D**). In some cases, populations found as two populations in one modality were represented as one in the other modality. For example, monocytes and were not separable from macrophage-derived TAM populations in IMC but were distinguishable by scRNAseq. Overall, these analyses allowed us to validate populations identified by IMC, and align them with their transcriptomic identities (**Fig 5C**)

**Figure 5.**
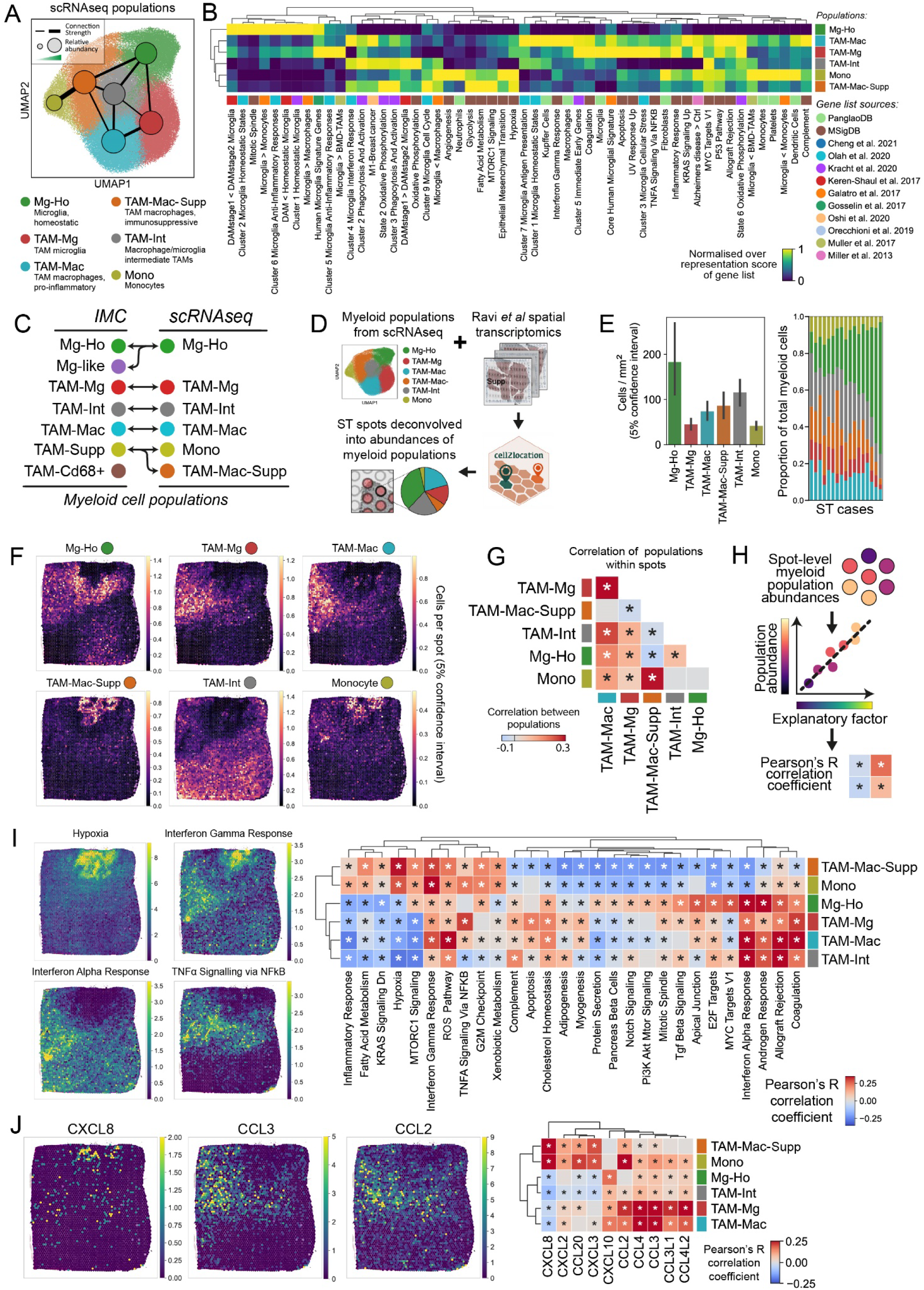
Spatial transcriptomics (ST) reveals hypoxia and chemokines as determinants of myeloid cell positioning in GBM. **A**. UMAP of the myeloid populations identified by Leiden clustering in the Ruiz *et al* single-cell sequencing (sc-RNA-seq) dataset. **B**. Comparison between the transcriptomes of the myeloid populations and published gene lists for biological processes (MSigDB)*(30)*, cell identities (PanglaoDB)*(31)*, myeloid cell phenotype in glioblastoma and other conditions disease *(7, 32–40)*. Enrichment of the gene lists was calculated by over representation analysis. **C**. Alignment of the populations identified by IMC and sc-RNA-seq. **D**. Schematic showing the strategy for deconvolving the Ravi *et al (41)* spatial transcriptomics datasets using the *cell2location (42)* and the transcriptomic signatures of the myeloid populations we identified in the Ruiz *et al* dataset *(29)*. **E**. Predicted abundances of the myeloid populations calculated using *cell2location*. **F**. Distribution of myeloid populations in ST spots within a representative deconvolved ST case. **G**. Pearson’s R correlation between the different myeloid population present in each spot. **H**. Strategy for identifying explanatory factors that may control the positioning of myeloid cell populations. Using this strategy, myeloid cell abundances were correlated with the transcriptomic signatures of biological processes from the MSigDB database (**I**), or expression of chemokines (**J**). *p<0.05, with correction for multiple Pearson’s R tests using the Benjamini-Hochberg procedure (**G, I, J**)

We then used the transcriptomic identities of the myeloid populations to deconvolve a spatial transcriptomic dataset of 19 GBM cases taken from tumour core (**Fig 5A**), allowing us to predict the abundances for each population in each case (**Fig5 D&E**). Spatially mapping the deconvolved populations to individual spots demonstrated compartmentalisation of populations, with clear segregation of where populations were found at their highest densities (**Fig 5F**). Where different populations were found in the same spot, positive correlations were found between populations with similar phenotypes (e.g. Mono and TAM-Mac-Supp), and negative correlations between dissimilar populations (e.g. Mg-Ho with TAM-Mac-Supp) (**Fig 5G**), supporting earlier observations made in IMC of phenotypically similar populations spatially associating (**Fig 2**).

To understand the potential drivers of myeloid positioning, we correlated the abundance of the myeloid cell populations with gene signatures of biological processes within each spot (**Fig 5H&I**). This confirmed findings made by IMC, with TAM-Mac-Supp and Mono populations preferentially accumulating in areas of increased hypoxic signalling and altered metabolism, which were associated with GLUT1 gene (SLC2A1) expression (**Supp 3E**). By contrast, the remaining myeloid populations were positively associated with signatures of interferon-α, androgen, and coagulation responses. Pro-inflammatory signalling pathways (interferon-γ, TNFα, and reactive oxygen species) also influenced the positioning of specific myeloid subpopulations. Repeating this analysis for chemokine genes suggested the positioning of specific subsets of myeloid cells were controlled by distinct groups of chemokines (**Fig 5J**). The same populations responsive to hypoxia (TAM-Mac-Supp and Mono) associated with CXCL8, CXCL2, CCL20, CXCL3, whereas TAM-Mg and TAM-Mac populations were associated with CCL4, CCL4, CCL3L1 and CCL4L2. By contrast, the positioning of Mg-Ho and TAM-Int were only weakly associated with chemokine expression. Together, these analyses reinforce and expand on findings made by IMC, showing that specific myeloid populations accumulate in hypoxic niches in the TME, and suggest a role of spatially variable chemokine– and inflammatory-signalling in myeloid compartmentalisation.

### Myeloid cell environments were defined by distinct patterns of myeloid populations within ST spots

Both IMC and ST analyses suggested that specific myeloid niches exist in the GBM tumour microenvironment, characterised by common biological processes (*e.g.* hypoxia), and occupied by myeloid populations with similar phenotypes. To identify these myeloid niches (hereafter termed myeloid environments), we clustered ST spots based upon their abundances of the 6 myeloid populations (**Fig 6A**). This identified five distinct myeloid environments, including three in which myeloid cells were highly abundant (0, 2 and 4) but in different combinations of phenotypically similar populations, and two with low abundance (1 and 3) of myeloid cells (**Fig 6B & C**). This supports observations made in IMC whereby if cells were clustered, it was with cells of the same or similar phenotype. Assessing the distribution of these environments within the TME found they organise into larger regions spanning several interconnected spots, with the proportion of connections between spots being therefore dominated by those between spots of the same environment (**Fig 6D**).

**Figure 6.**
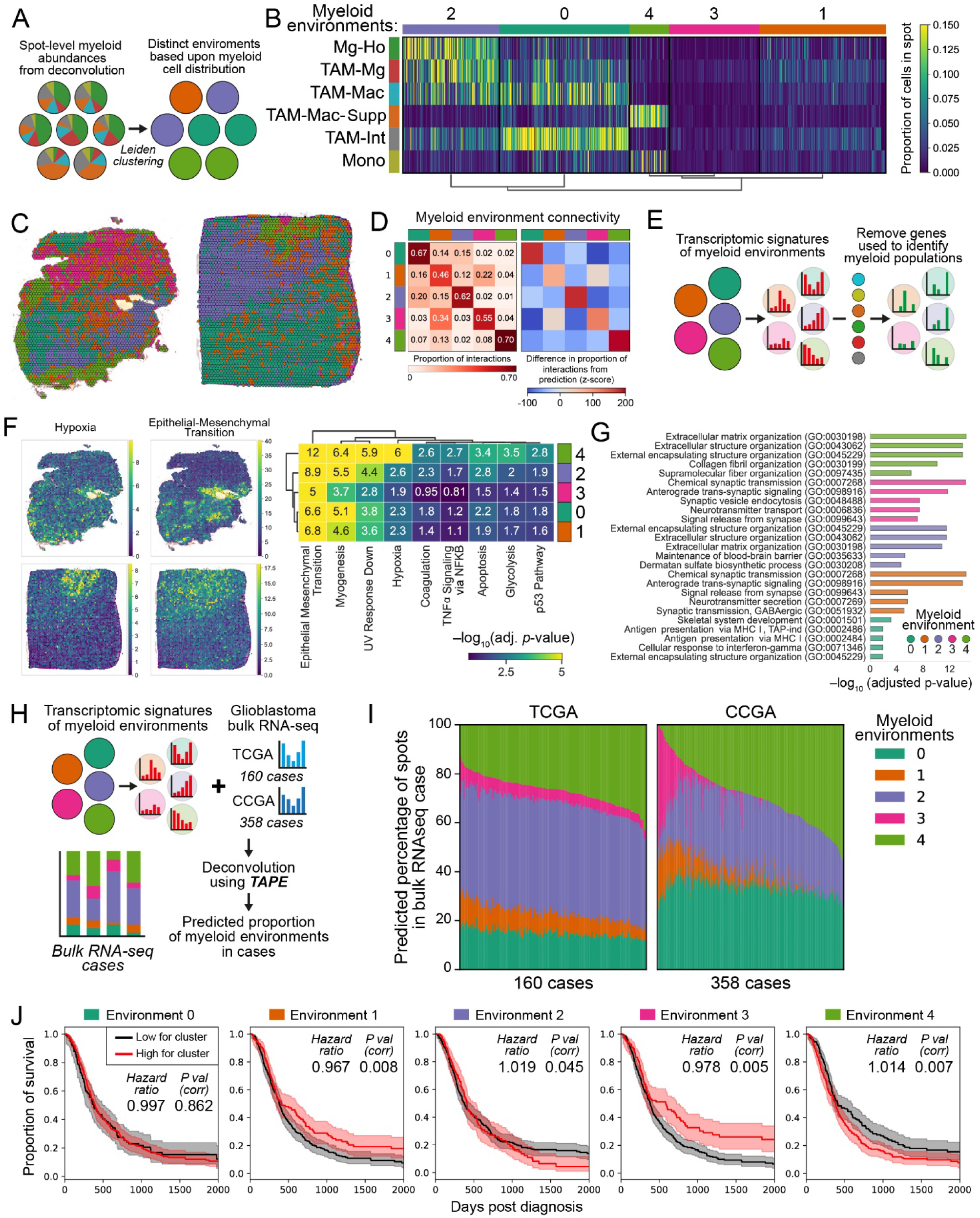
– Spatial clustering of myeloid cells is associated with poor outcome in glioblastoma. **A**. Strategy to identify environments in the deconvolved ST cases that have distinct patterns of myeloid cell distribution. **B**. Identification of distinct patterns of myeloid cell distribution using *k*-means clustering, corresponding to 5 distinct myeloid cell environments. **C**. Mapping of the 5 myeloid environments in example ST cases. **D**. Proportion of interactions between spots from each myeloid environment, assuming each spot is connected to its neighbouring 6 spots. **E**. Strategy to identify the transcriptomic changes arising from the non-myeloid cells in each spot, see Methods. **F**. Over representation analysis of the signatures of biological process from the MSigDB database *(30)* in the remaining non-myeloid genes in the different myeloid environments. **G**. The top 5 enriched terms from the Gene Ontology database in the non-myeloid genes in the different myeloid environments. **H**. Strategy to deconvolve bulk RNA sequencing GBM cases from the TCGA (The Cancer Genome Atlas) and CCGA (Chinese Glioma Genome Atlas) using the *TAPE* algorithm *(43–45)*, therefore allowing us to predict the proportions of myeloid environments in each case. **I**. Proportion of myeloid environments predicted by the TAPE algorithm in the TCGA and CCGA cases. **J**. Modelling the relationship between the abundance of each myeloid environment and glioblastoma survival using Cox proportional-hazards, with correction for multiple tests using Holm-Šídák. Hazard ratios are increase in risk of death per percentage point of myeloid environment abundance. Kaplan-Meier curves compare patients having the high (top 50%) or low estimates for the presence of that environment, shaded areas are 95% confidence intervals.

### Contribution of the non-myeloid compartment to the myeloid environments

We then analysed how the non-myeloid compartment varied between the different myeloid environments, specifically focusing on genes not differentially expressed between myeloid populations (**Fig 6E**). In this non-myeloid compartment, there was a clear signature of hypoxia and metabolism in environment 4 (**Fig 6F**), an environment enriched for myeloid populations which individually correlated with the signature of hypoxia (**Fig 5I**). This supports the existence of hypoxic niches in the tissue which affect the behaviour and positioning of both myeloid and tumour cells. In all 5 environments there were processes likely arising from the tumour compartment (*e.g.* epithelial-mesenchymal transition, p53 pathway). To obtain further insight into the non-myeloid factors which differentiate the myeloid environments, we performed gene ontology on the non-myeloid differentially expressed genes between myeloid environments (**Supp 4A, Fig 5G**). This demonstrated that environments 2 and 4 each had unique differences in extracellular matrix composition. Given these two environments were enriched for phenotypically different myeloid cells, it suggests that myeloid cell compartmentalisation may be influenced by the local deposition of extracellular matrix components by non-myeloid cells. The two environments with lowest abundance for myeloid cells (1 and 3), were enriched for genes associated with neuronal functioning, suggesting that neural-progenitor-like tumour cells may influence myeloid positioning by inhibiting local myeloid recruitment.

### The transcriptomic signature of specific myeloid environments was associated with reduced disease survival

The complete transcriptomic signatures of the myeloid environments were then used to deconvolve 518 bulk mRNA sequencing GBM cases from the TCGA and CCGA datasets using the TAPE algorithm (**Fig 6H**) *(43–45)*. This analysis allowed us to predict the proportions of myeloid environments present in each case, showing clear variability between patients, particularly in environment 4 which is dominated by TAM-Mac-Supp and monocytes (**Fig 6I**). Comparing the overall survival curves for patients either low or high for each myeloid environment (either lower or higher than the mean) and performing univariate COX proportional hazard analyses showed that the abundance of the myeloid environments has a significant effect on GBM survival. Specifically, tumours with high proportions of environment 2 (featuring clustering of pro-inflammatory populations) and 4 (featuring clustering of immunosuppressive populations) were associated with worse survival, whereas high proportions of environments 1 and 3 (with low myeloid clustering) was associated with better survival. As the proportions were correlated with one another, a multivariate model with the 5 environments could not be built. However, a multivariate model using principal components of the 5 environments also showed a significant effect of varying myeloid abundance on survival (**Supp 4B**). Together, these IMC and spatial transcriptomics analyses demonstrate that the spatially regulated compartmentalisation of myeloid cell populations, which is instructed through tumour environmental cues, contributes to disease trajectory during GBM.

## Discussion

In this study we have revealed the compartmentalisation of key myeloid cell populations within GBM. Utilising orthogonal high dimensional IMC analyses and deconvolved ST datasets, we robustly identified at least six myeloid cell populations. Microglia existed on a spectrum of activation states from homeostatic to pro-inflammatory activation, with the transition being associated with a reduction of P2RY12, supporting previous observations made in GBM *(7, 19, 25)*. Macrophages were either immunosuppressive, showing upregulation of markers previously associated with an immunosuppressive phenotype in GBM myeloid cells (CD163, CD206) *(11, 17, 19, 20)*, or proinflammatory. In IMC analyses, these pro-inflammatory macrophages were distinguishable from pro-inflammatory microglia by further reduction in P2RY12 expression, and upregulation of CD14 and CD16. Furthermore, transcriptomic analysis demonstrated they had increased interferon-γ and complement signalling. Although we could identify these myeloid states as distinct populations, there was usually a gradient change in marker expression between populations which suggested cells were transitioning between phenotypes. Specifically, diffusion pseudo time analysis suggested microglia undergo pro-inflammatory activation, adopt a more macrophage-like phenotype, before finally becoming immunosuppressive. This is in line with previous studies in GBM, which report myeloid cells in transitionary states between more obviously identifiable pro-inflammatory or suppressive phenotypes *(4, 21, 22)*, with cells able to simultaneously co-express M1– and M2-markers *(7)*. Consequently, in both IMC and sequencing datasets we found canonical strongly polarised TAM and microglial populations, as well as large intermediate populations that exhibited characteristics of microglia and macrophages, and of mixed pro– and anti-inflammatory polarisation.

Our IMC analyses demonstrated that microglial cells tended to localise within the GBM tumour edge and macrophage-derived TAMs to accumulate in the tumour core, supporting previous observations made in both human GBM and in murine models *(11, 15, 21, 24, 46, 47)*. Importantly, however, this compartmentalisation was not mutually exclusive, as the numerous myeloid populations were present in each tumour region analysed. However, the positioning of the different myeloid cells within the TME was non-random: myeloid cells preferentially positioned with cells of the same or similar phenotype, and avoided interacting with myeloid cells of a dissimilar phenotype. This behaviour differed in strength between populations, being strongest in immunosuppressive TAMs, and was also recapitulated when analysing ST datasets. Furthermore, most cell-to-cell interactions made by myeloid cells were with other myeloid cells. These analyses suggest myeloid cell interactions as a major cellular determinant of myeloid cell positioning, which is not unexpected given their role as the primary producers of chemokines and other signals that elicit responses from myeloid cells. Notably, although the abundance of certain myeloid populations differed between the edge and core regions of the tumour, the overall compartmentalization behaviour of myeloid populations was broadly similar in both parts of the tumour. This suggests that the factors that coordinate myeloid cell activities (whether deriving from myeloid or tumour cells, or the environment) are conserved throughout the GBM TME.

Hypoxia was identified as a major controller of myeloid cell compartmentalisation within GBM, particularly influencing immunosuppressive populations. In IMC analyses, immunosuppressive myeloid cells (CD163+, CD206+) were found to be most strongly linked to hypoxic areas, although this association was also observed, albeit less significantly, in other macrophage populations. In contrast, non-activated microglia exhibited the lowest association with hypoxia across all analyses. This is in support of previous studies that have reported that macrophages, rather than microglia, accumulate in hypoxic areas of the TME *(9, 19)*. Immunosuppressive myeloid cells also showed an interaction with the vasculature in IMC analyses that was consistently observed throughout the TME. Further examination of this population by transcriptomics revealed it was a combination of monocytes and immunosuppressive macrophages, explaining their vascular association. In agreement with IMC analyses, both monocytes and immunosuppressive macrophages exhibited a preferential localization within hypoxic niches in ST analyses. However, the presence of hypoxic signalling within the monocytes implied that most were already being influenced by the GBM environment and were likely differentiating into macrophages. Indeed, we saw greater infiltration of immunosuppressive myeloid cells in IMC analyses into hypoxic areas of the tumour core. These observations are supported by previous characterisations of immunosuppressive myeloid cells in GBM, which have typically found them to be blood-derived macrophages with high expression of hypoxia-related genes *(9, 16, 17, 24)*. Together, our findings from IMC, ST, and scRNAseq all indicate that hypoxia plays a pivotal role in the differentiation of myeloid populations in GBM towards an immunosuppressive phenotype, as reported in TAMs in various cancer types *(48)*.

Accumulation of myeloid cells in hypoxic niches could also be attributed to the release of chemotactic signals induced by hypoxia *(48)*. Indeed, our investigation unveiled heterogeneous expression of chemokines throughout the TME, with specific chemokines correlating with immunosuppressive populations in hypoxic regions. Other chemokines (e.g. CCL-2, 3 and 4) were associated with the positioning of pro-inflammatory macrophage and microglial populations. This in agreement with previous studies reporting similar populations located at the tumour periphery serve as the primary source of these chemokines, with authors hypothesising they are responsible for recruiting additional myeloid cells *(5, 24, 46)*. Alternatively, these chemokines may themselves affect myeloid phenotype, with CCR2 knock out resulting in a greater proportion of microglia in mouse tumours, though not by blocking monocyte infiltration, but by blocking monocyte-to-macrophage differentiation *(19)*.

It is also highly probable that myeloid compartmentalization is influenced by the positioning and interaction with tumour cells. However, compared to some solid tumours, the different components of the TME in GBM do not commonly show a macroscopically obvious segregation into defined immune and tumour compartments, likely due to the highly infiltrative nature of tumour cells in glioblastoma. Surprisingly, we found that all myeloid populations showed some degree of spatial avoidance of tumour and neuroglial cells in the TME. A potential caveat of our analyses is that we did not differentiate between the three known neoplastic subtypes present in GBM *(23, 49)*. However, existing data suggests they may differentially shape the myeloid landscapes in GBM. For example, GBM tumours rich in mesenchymal subtype have the highest myeloid cell density, while those abundant in pro-neural subtype show the lowest macrophage proportion *(6, 50)*. Myeloid cells can directly shape tumour cell fate, with macrophages being shown to drive differentiation of tumour cells towards a mesenchymal phenotype *(51)*. Further studies are therefore required to address how specific myeloid and neoplastic cell subtypes interact during GBM.

The clinical significance of the spatial arrangement of immune cells within a tumour has been demonstrated in various cancer types *(20, 52, 53)*, frequently offering superior prognostic value compared to the mere abundance of immune cells *(54–56)*. Due to the previously described clustering of myeloid populations in different areas of the TME, we were able to extract the transcriptomic signatures of the different spatial arrangements of myeloid populations. These different myeloid environments were each dominated by myeloid cells of a different phenotype. When we then deconvolved 518 bulk RNAseq cases using these signatures, we found reduced survival time in patients with tumours that were enriched with environments where either proinflammatory or immunosuppressive populations were highly clustered. This demonstrates, for the first time, that the topology of myeloid populations in GBM is associated with disease outcome. Specific immunosuppressive myeloid populations identified using sc-RNA seq (e.g. CD73 and MARCO high) have been associated with poor survival in GBM *(9, 10)*. Our spatial analyses add context to these findings, suggesting these cells are located in hypoxic areas, and are likely vascular-associated. Pro-inflammatory macrophages are associated with disease progression in lower grade gliomas, with fewer immunosuppressive macrophages compared to GBM *(57–59)*. The detrimental effect of proinflammatory niches may therefore represent areas transforming from low to high-grade tumour.

Overall, these results align with an expanding body of research that indicates that the spatial structure of the GBM TME plays a decisive role in determining the clinical course *(20, 60)*. Ultimately, better understanding of the biology of GBM and revealing how cells communicate within the TME will allow us to start deconstructing the heterogeneity in GBM, and stratify patients for targeted treatments.

## Materials and Methods

### Clinical samples for imaging mass cytometry

Eight primary IDH^wt^ glioblastoma cases were retrieved from the Department of Cellular Pathology at Salford Royal Hospital bank (**Table 1**, Ethics IRAS ID 244538). The interface between tumour and cortex (edge), and the tumour core were annotated by a neuropathologist on H&E stained sections. Tissue micro arrays (TMAs) were subsequently generated using 3 mm diameter cores, with 3 cores taken per case.

### Imaging mass cytometry tissue staining

Sections from tissue micro arrays (5 µm thickness) underwent staining with lanthanide-conjugated antibodies as instructed by manufacturer *(61)*. In brief, sections underwent deparaffinisation, followed by antigen retrieval at 96°C for 30 min in Tris-EDTA at pH 8.5. Non-specific binding was blocked with 3% bovine serum albumin for 45 min, followed by incubation with lanthanide-conjugated primary antibodies (overnight at 4°C) which were diluted in PBS with 0.5% BSA (**Table 2**). Antibodies were conjugated with metals using Maxpar Antibody Labeling Kits (Standard BioTools) and were validated with positive control tissue (tonsil for immune-targeted antibodies) and dilutions optimised with GBM tissue. Slides were then washed with PBS and 0.1% Triton-X100 in PBS. Slides then underwent nuclear staining with iridium (1:400, Intercalator-Ir, Standard Bio Tools) for 30 minutes at RT, before being briefly (10 s) washed with ultrapure water and air-dried. Images were acquired of metal-stained tissue sections on a Hyperion imaging mass cytometer as per manufacturer’s instructions (Standard BioTools). Each TMA core was imaged in a separate region of interest. In brief, the tissue was laser-ablated in a rastered pattern in a series of 1 µm^2^ pixels. The resulting plume of ablated tissue was then passed through a plasma source, ionising it completely into its constituent atoms. Time-of-flight mass spectrometry then discriminated the signal for each of the metal-conjugated antibodies, and images for each antibody were reconstructed based off the metal abundancy at each pixel. Staining was reviewed by a neuropathologist using MCD Viewer (Standard BioTools). In representative images of IMC data shot noise was removed using the IMC-Denoise algorithm *(62)*.

**Table 2.** Antibody panel for imaging mass cytometry.

| Metal channel | Antigen | Clone / Catalogue # | Supplier |
| --- | --- | --- | --- |
| 89 | Smooth muscle actin | 1A4 | Abcam |
| 113 | Cd68 | KP1 | BioLegend |
| 115 | Cd235ab | KIR2 | BioLegend |
| 139 | Pan-cytokeratin | AE-1/AE-3 | BioLegend |
| 141 | S100B | EP1576Y | Abcam |
| 142 | MHC1 | EMR8-5 | Abcam |
| 143 | Vimentin | RV202 | Standard BioTools |
| 144 | Cd14 | D7A2T | Cell Signalling |
| 145 | Ki67 | B56 | BD Pharmingen |
| 146 | Cd16 | SP175 | Abcam |
| 148 | Cd66b | G10F5 | Novus |
| 149 | Cd11b | EP1345Y | Abcam |
| 150 | Cd44 | IM7 | BioLegend |
| 151 | Granzyme B | EPR20129-217 | Abcam |
| 152 | Cd45 | CD45-2B11 | eBioscience |
| 153 | Cd31 | JC/70A | Novus |
| 154 | Cd11c | EP1347Y | Abcam |
| 155 | HIF1a | EP12154 | Abcam |
| 156 | Cd4 | EPR6855 | Abcam |
| 158 | Cd109 | C9 | Santa Cruz |
| 159 | Olig2 | Polyclonal, AB9610 | Millipore |
| 160 | Vista | D5L5T | Cell Signalling |
| 161 | iba1 | Polyclonal, 019-19741 | WAKO |
| 162 | Cd8a | CD8/144B | eBioscience |
| 163 | GLUT1 | EPR3915 | Abcam |
| 164 | Nestin | 25/Nestin | Biologend |
| 165 | Fibrinogen | EPR18145-84 | Abcam |
| 166 | Cd74 | LN2 | Biologend |
| 167 | Met | Met | D1C2 |
| 168 | P2RY12 | HPA014518-100UL | Merck |
| 169 | Cd163 | EDHu-1 | BioRad |
| 170 | Cd3 | D7A6E | Cell Signalling |
| 171 | pERK1/2 | D13.14.4E | Cell Signalling |
| 172 | TMEM119 | Polyclonal, ab185333 | Abcam |
| 173 | Sox2 | 245610 | R&D |
| 174 | MHCII | TAL1B5 | Abcam |
| 175 | Cd206 | E2L9N | Cell Signalling |
| 176 | GFAP | GA5 | Sigma |

### Cell segmentation of IMC images

Single-cell information was extracted from IMC images using an established protocol *(63)*. In brief, stacks of TIFF images were extracted from MCD files for each region of interest whereby individual channels corresponded to each lathanide-conjugated antibody. Ilastik *(64)* was then used to produce a pixel probability classifier that identified background, cytoplasmic and nuclear pixels. The resulting pixel probability maps were then converted into cell segmentation masks that identified the regions corresponding to individual cell boundaries. These cell segmentation masks were then applied to each of the antibody channels, generating single-cell expression data for each of the channels, along with the spatial context of where the cell was in the tissue.

### Analysis of single-cell IMC data

Single-cell IMC data was analysed in Python using packages designed to analyse single-cell data (Scanpy, v1.9.3), and spatial molecular data (Squidpy v1.2.3, and ATHENA v0.1.3) *(65–67)*. The mean cell intensity of each marker was normalised to the 99^th^ percentile of its expression. Leiden clustering *(18)* was then used to identify cell populations present in the IMC data, which were then manually annotated based upon patterns of marker expression corresponding to known cell types and activation states. The transition between myeloid populations was assessed using diffusion pseudotime and PAGA analyses *(68)* using Scanpy.

### Spatial distribution and interaction analyses in IMC data

Metrics were calculated at the single-cell level, before being mean averaged at the population level for each region of interest (ROI). For spatial distribution analyses, the distance between each cell and the nearest member of each other population was calculated. For interaction analyses, the number of interactions made by each cell (either calculated as 6-nearest-neighbours or cell-to-cell contact) to other populations was calculated and expressed as a proportion of total interactions. The observed values for mean distances and proportions of interactions were then subtracted from the values predicted by a random distribution of cells, which was calculated by randomly distributing cell labels 300 times within each region. This corrected interactions for differences between regions in abundances of interacting populations. The resulting differences between observed and predicted values were separately averaged across all edge and core regions, with statistical difference from random distribution (observed – predicted = 0) assessed using Wilcoxon tests with Benjamini-Hochberg correction. Rao’s quadratic entropy is a measure of phenotypic heterogeneity and was calculated between each cell and the cells it makes direct cell-to-cell contact using ATHENA *(67)*, with values mean averaged at the population levels for each region.

### Clustering and assortativity measures of myeloid cells in IMC data

Clustering coefficients *(26)* and assortativity *(27)* were calculated on 6-nearest-neighbour graph of myeloid cells in each region using Networkx (v3.1). A high clustering coefficient designates a population forms densely interconnected clusters, and a low value suggests that population is more loosely clustered in the TME *(26)*. Assortativity is another measure of clustering in a network, and measures the tendency of cells to connect to cells of the same population, rather than cells of a different population *(27)*. Clustering coefficients for each population were calculated by extracting each population as a subgraph and calculating their average clustering coefficients. The observed clustering coefficients were then compared to a random distribution as described above, whereby cell labels were randomly distributed 300 times.

### Cell environment analysis in IMC data

The environmental expression of GLUT1, pERK1/2 and fibrinogen was calculated by taking the mean average expression of each marker in a 40 µm diameter window centred on each cell. The distribution of the resulting environments was then statistically compared between cells from either edge or core of the tumour or between different populations using linear mixed models in which cells were nested within regions, which were nested within individual cases. Correction for tests was performed using a Holk-Šídák correction.

### Re-analysis of myeloid cells from a single-cell sequencing dataset

We analysed 127,339 myeloid cells from a multi-study single-cell RNA sequencing dataset that incorporated 240 patients from 26 separate sequencing studies *(29)*. We analysed a 25% random sample of cells labelled as microglia, macrophages or monocytes by the original authors. Batch effects were corrected using the Harmony algorithm *(69)*, populations identified using Leiden clustering *(18)*, and connectivity of the populations assessed using PAGA analysis *(66, 68)*. Over expression analyses were performed with the Decouplr (v1.3.4) using hypergeometric tests (false discovery rate < 0.05). These used published gene lists from studies assessing the phenotypes of microglial in other conditions *(32–37)*, of myeloid cell activation *(38)*, and myeloid cells in GBM *(7)* and other cancers *(39, 40)*. Gene lists from canonical pathways of biological processes provided by MSigDB *(30)*, and of cellular identities from PanglaoDB *(31)* were also used. Activity inference for pathways from the PROGENy database *(70)* were performed using multivariate linear modelling.

### Deconvolution of spatial transcriptomics dataset

We analysed a published spatial transcriptomic (ST) dataset of 19 IDH^WT^ GBM cases taken from tumour core *(41)*. Data analysis was performed using Scanpy. In brief, low-quality spots (<1000 counts) and mitochondrial genes were removed, counts were normalised per cell, and log transformed. The cell-type composition of each spot was then calculated using cell2location (v0.1.3) *(42)*. Reference expression signatures of the myeloid cell populations were created from the single-cell sequencing dataset by taking the mean over all cells within each population. Reference signatures for non-myeloid populations were similarly created from the ‘*annotation level 3*’ labels from the Ruiz *et al* dataset *(29)*, and were included in the matrix of reference expression signatures to account for all potential cell populations present in the GBM TME, ensuring accurate deconvolution. The resulting abundances are the lower limit at which the model is confident, in other words, at least this amount is present.

### Predicting transcriptomic controllers of myeloid cell positioning

To understand factors that control myeloid cell positioning, the abundance of each myeloid cell population estimated from deconvolution was correlated with potential explanatory factors using Pearson’s R correlation. Factors that did not vary between populations (<0.05 STD) were removed, and remaining analyses were corrected for multiple tests using a Benjamini-Hochberg correction. Explanatory factors were either single genes (chemokines), or signatures of hallmark biological processes provided by MSigDB *(30)* which were quantified in each spot using hypergeometric tests in Decouplr.

### Identification of myeloid environments and their transcriptomic signatures

To identify the different myeloid environments, spots were clustered based upon the estimated abundance of myeloid cells populations from deconvolution. Population data was scaled to unit variance and zero mean, and batch corrected between cases using the BBKNN algorithm *(71)*. Distinct patterns of myeloid abundance (constituting different environments) were identified using Leiden clustering in Scanpy *(18)*. For assessments of connectivity between environments, each spot was connected to each of its surrounding 6 spots. To investigate the contribution of non-myeloid cells to the transcriptomes of the different environments, any genes with >0.1 (counts) STD between myeloid populations in the reference expression signatures used for deconvolution of ST data were excluded from analysis (2115 genes removed, leaving 10031. The remaining genes were then compared between environments using hypergeometric tests using gene lists from MSigDB, and by calculating differentially expressed genes (DEGs) using Wilcoxon rank-sum test *(72)*. The resulting DEGs (*p*<0.01, 1.5 fold enrichment, 117 genes per cluster) were then used for over-representation analysis by Enrichr web services via GSEApy (v1.0.4), accessing the Gene Ontology databases *(30, 73, 74)*.

### Deconvolution of bulk sequencing

The transcriptomic signatures of the myeloid environments were used to deconvolve bulk mRNA sequenced from IDH^wt^ GBM cases from the TCGA PanCancer atlas (160 cases) and CCGA (358 databases) *(44, 45)*. Bulk mRNA data from both datasets were independently sequenced on the Illumina HiSeq V2 platform, count data RSEM (RNA-Seq by Expectation Maximization) normalized, and batch normalized *(44, 45)*. Deconvolution to estimate the proportion of myeloid environments in each bulk case was then performed using the TAPE algorithm *(43)*, which sampled 500 spots from each myeloid environment, and was ran with the following hyperparameters: variance threshold 0.99, min-max scaling. The resulting proportions of myeloid environments were associated to patient survival using Cox proportional hazards models ran using the ehrapy (v0.3.0) *(75)*. Kaplan-Meier curves compare patients having the high (top 50%) or low estimates for the presence of that environment.

### Statistical analyses

Statistical tests were performed using the statsmodels (v0.13.5) and ehrapy packages, and are specified for individual methods. For cell environment analyses, calculations were at the cell-level, and so for linear mixed models (LMMs) both region and case were used as grouping factors. For all other LMMs, data was mean averaged at the region level, and patient case was used a grouping factor. Where LMMs were used concurrently, a Holm-Šídák correction was used for calculation of *p* values.

## Acknowledgments

The authors would like to thank the staff of the University of Manchester Flow Cytometry and Bioimaging core facilities for their assistance, and the staff of Salford Royal Hospital pathology department.

## Author contributions

Conceptualization: MJH, KC, LB

Methodology: MJH, SG, NG

Investigation: MJH, LB, GH

Resources: JM, FR

Software: SG, NG Visualisation: MJH

Supervision: KC, DB, FR, OP, DC, AK, DW, SA,

Writing – original draft: MJH, KC

Writing – review & editing: MJH, DC, FR, AK, DW, SA, OP, DB, KC

Funding acquisition: KC, DB, OP, SA, AK

## Competing interests

Authors declare that they have no competing interests.

## Data and materials availability

Single-cell sequencing and spatial transcriptomics datasets are accessible from their original publications *(29, 41)*. IMC data (TIFF images and single-cell data) is retrievable from the following repository: doi:10.5061/dryad.2z34tmprw.

---

**Supplemental 1.**
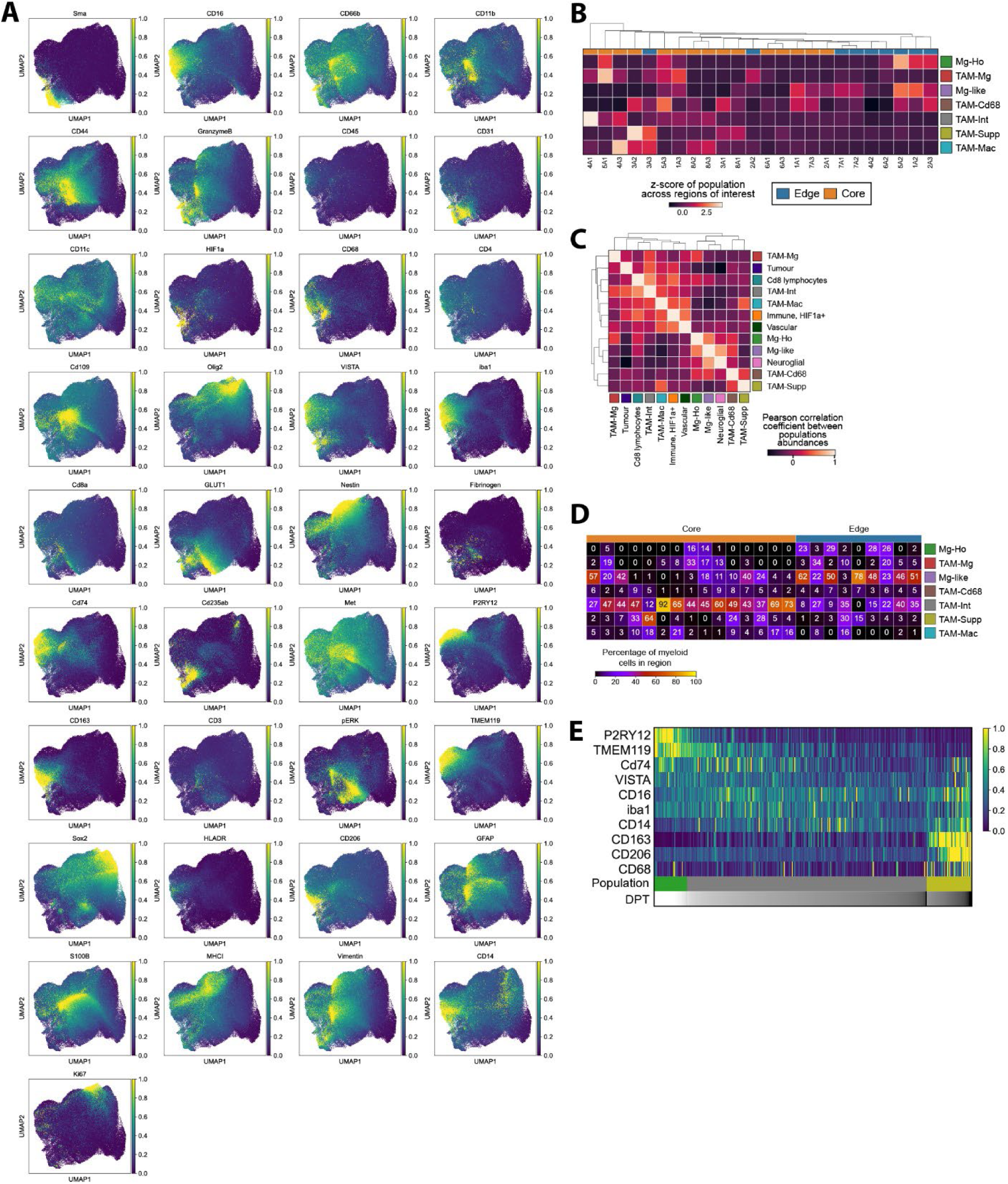
**A**. UMAPs visualising the single-cell data acquired from the IMC workflow from all cases, demonstrating the distribution of all the markers in the panel. Each marker is normalised to the 99^th^ percentile of its expression. **B**. *K*-means clustering of regions of interest based upon myeloid cell abundance. **C**. Pearson’s correlation coefficient between populations in each region. **D**. Percentage of total myeloid cells made up by each population in each region. **E**. Diffusion pseudotime modelling a potential pathway of differentiation from homeostatic microglia (Mg-Ho), through a macrophage/microglia intermediary state (TAM-Int), into immunosuppressive myeloid cells (TAM-Supp).

**Supplemental 2.**
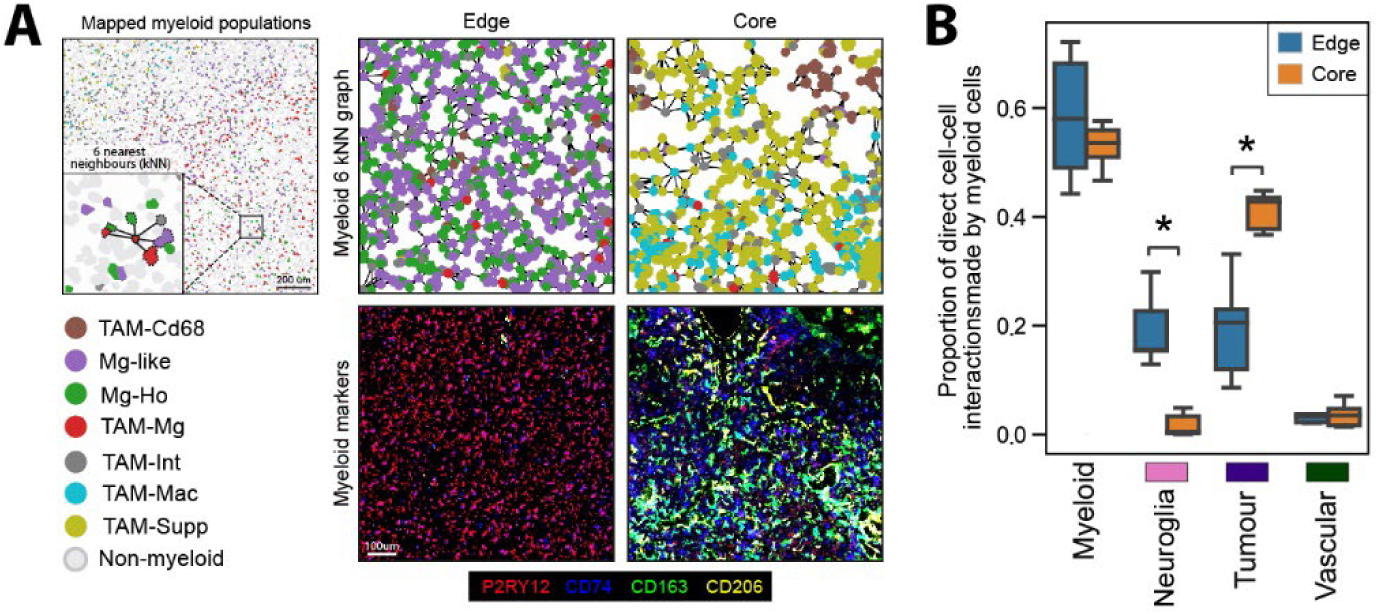
**A**. Demonstration of how the distribution of myeloid cells can be assessed by connecting each myeloid cell to its nearest 6 neighbours, with representative examples of the resulting graphs from edge and core regions, and associated IMC images for myeloid markers. **B**. Proportion of cell-cell interactions made by all myeloid cells (not broken down into separate populations) with non-myeloid cells. Comparison made by linear mixed model.

**Supplemental 3.**
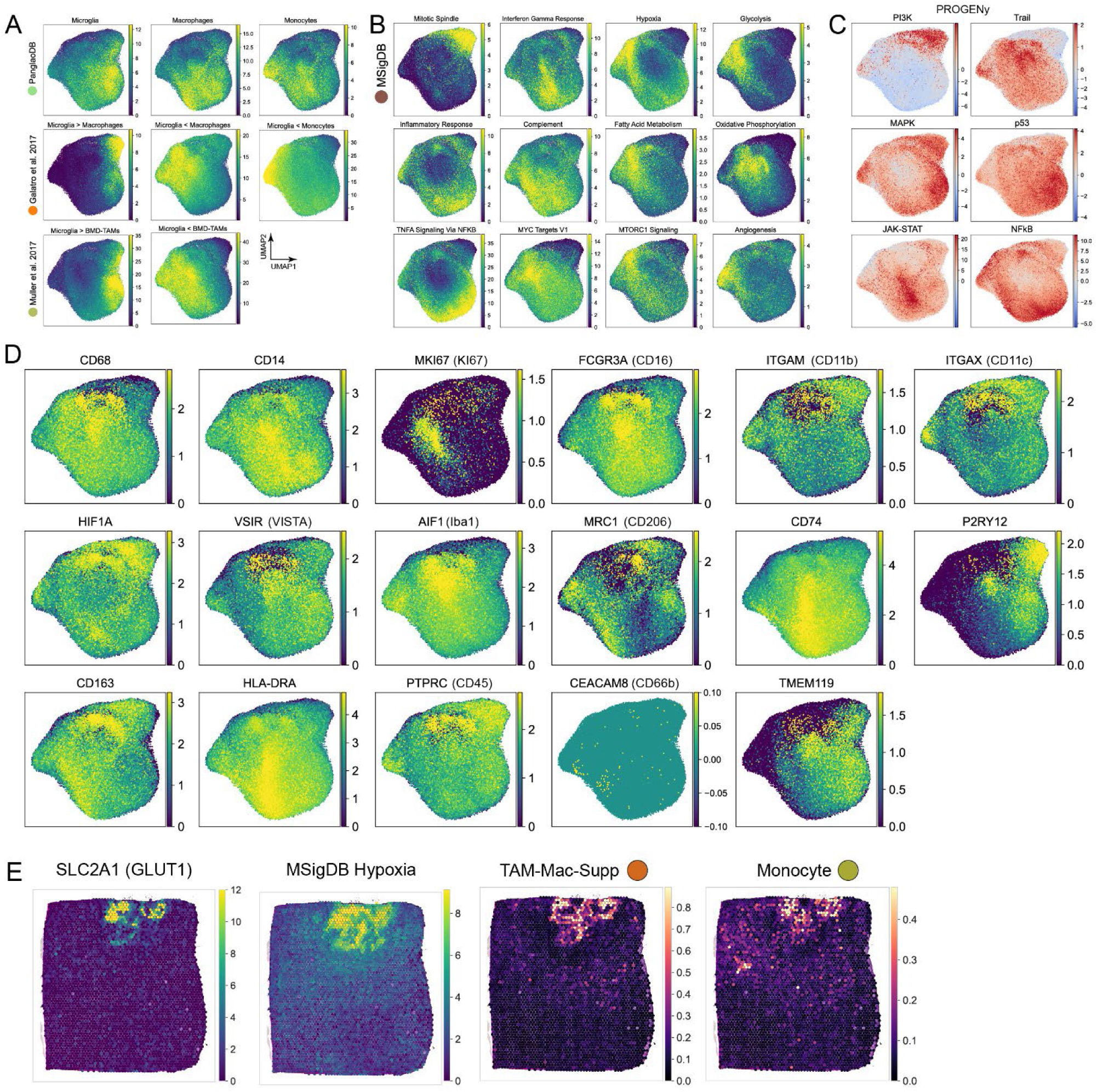
Enrichment of specified gene lists was calculated by over representation analysis (ORA) for individual cells, which was then plotted by UMAP. **A**. ORA of gene lists differentiating microglia, macrophages, and monocytes *(6, 55, 62)*. **B**. ORA of biological processes from the MSigDB database *(61)*. **C**. Activation or suppression of signalling pathways assessed by reference to the PROGENy database *(63)*. These analyses (**A, B, C**) were performed using the *Decouplr* package in Python. **D**. Expression of the genes encoding the myeloid proteins targeted by the IMC antibody panel. **E**. Representative ST case showing the distribution of the GLUT1 gene (SLC2A1) alongside the MSigDB hypoxia signature, and abundance of immunosuppressive macrophages (TAM-Mac-Supp) and monocytes.

**Supplemental 4.**
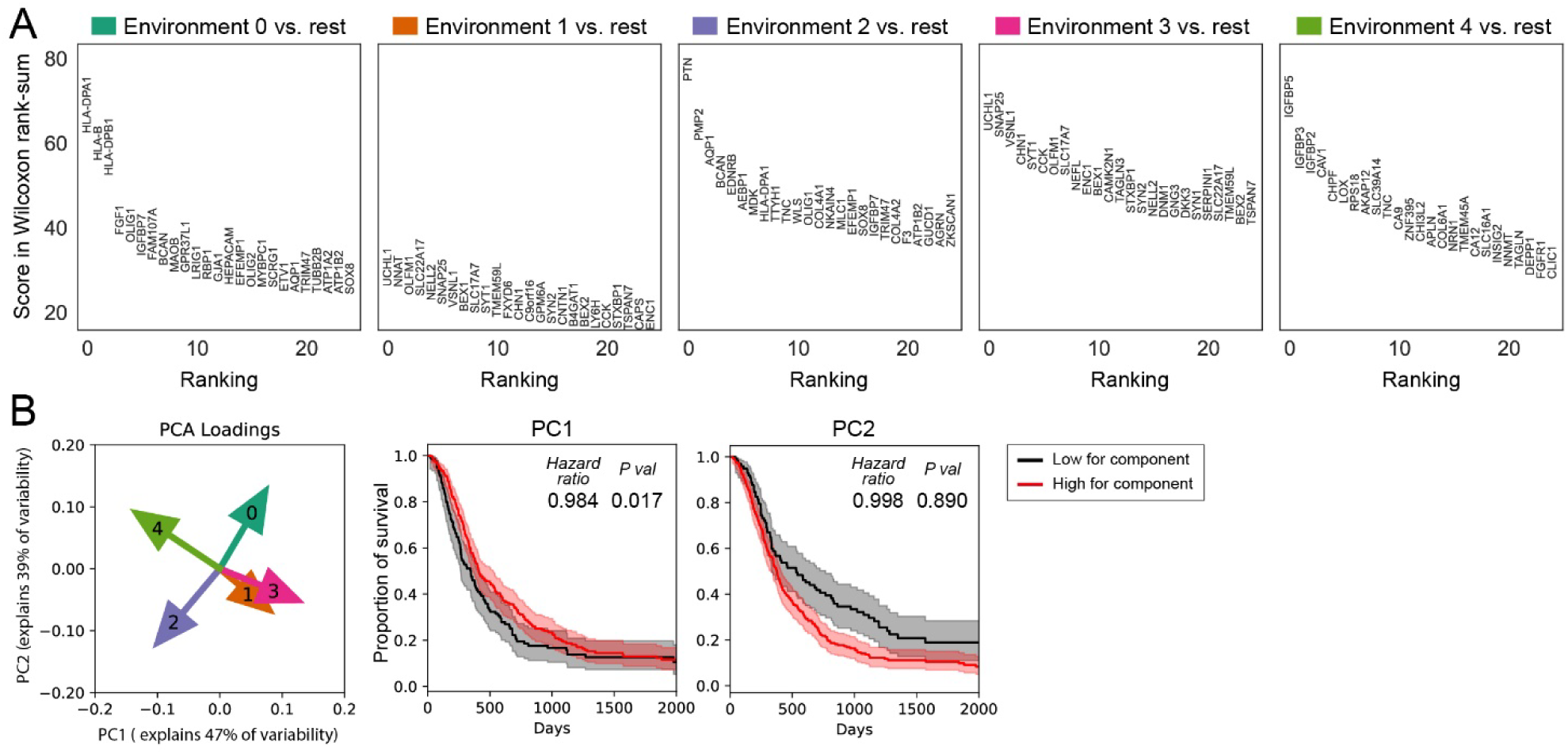
**A**. The genetic changes associated with non-myeloid cells in each myeloid environment were extracted as described in **Figure 6E**. The top 25 differentially expressed non-myeloid genes differentiating each myeloid environment from the others was then calculated by Wilcoxon rank-sum analysis. **B**. Principal components analysis was used to summarise the abundance of the 5 myeloid environments into two principal components, which are by definition not correlated. The relationship between these principal components and GBM survival was modelled using Cox proportional-hazards, and top 50% and bottom 50% of patients for these PCs compared by Kaplan-Meier curves, shaded areas indicate 95% confidence intervals.

## Notes

### Competing Interest Statement

The authors have declared no competing interest.

